# The Phosphorylation State of Melanophilin, a Myosin Va Adapter Protein, Biases the Tug-of-War Between Myosin Va and Kinesin-1 on Liposomes at Actin-Microtubule Intersections *In Vitro*

**DOI:** 10.64898/2026.09.01.748553

**Authors:** Brandon M. Bensel, Patricia M. Fagnant, Jill E. McFarlane, Oban Galbraith, Michael J. Previs, Kathleen M. Trybus, David M. Warshaw

**Affiliations:** Department of Molecular Physiology and Biophysics, University of Vermont Larner College of Medicine, Burlington, VT 05405

## Abstract

In cells, myosin Va and kinesin-1 motors navigate their shared vesicular cargos through numerous actin-microtubule (MT) intersections. How cargos exit these intersections to ensure precise destination delivery remains unclear. We hypothesized that the myosin Va adapter melanophilin – a phosphorylated protein that links myosin Va to melanosomes but can also bind actin and MTs – acts as a phosphorylation-dependent switch to bias the tug-of-war between cargo-bound myosin Va and kinesin-1 at actin-MT intersections. We modeled melanosome transport *in vitro* using 350-nm liposomes with ∼5 each of surface-bound constitutively active myosin Va and kinesin-1, and ∼5 full-length melanophilin with varying phosphorylation levels. Regardless of the track the liposome entered the actin-MT intersection on, phosphorylated melanophilin biased liposomes to exit on actin, whereas dephosphorylated melanophilin biased liposomes to exit on MTs. Consistent with this, phosphorylated melanophilin preferentially bound actin 2-fold over MTs, and slowed myosin Va liposome transport on actin by ∼40%, effectively acting as an anchor. Conversely, dephosphorylated melanophilin preferentially bound (2-fold) MTs over actin, and increased kinesin-1 liposome transport distance on MTs by acting as a tether. Therefore, melanophilin, depending on its phosphorylation state, biases track selection by kinesin-1/myosin Va transported cargos in actin-MT intersections that exist in complex cellular cytoskeletal networks.

**Significance Statement:**

- Cargos transported by myosin Va and kinesin-1 motors must select an exit track at actin-MT intersections. Melanophilin, a myosin Va adapter that attaches myosin to melanosomes, has phosphorylation-dependent actin or MT binding preference, potentially biasing the tug-of-war between cargo-bound kinesin-1 and myosin Va in actin-MT intersections.
- Physiologically relevant liposomes with surface-bound myosin Va, kinesin-1, and phosphorylated melanophilin preferentially exit actin-MT intersections on actin while liposomes with dephosphorylated melanophilin preferentially exit on MTs.
- These results support a novel track selection mechanism whereby adapter proteins on cargo bias the tug-of-war between distinct motors through phosphorylation-dependent track binding interactions.

## Introduction

Secretory vesicle transport involves microtubule- and actin-based molecular motors maneuvering their shared cargo through the cell’s complex cytoskeletal network (Fig. 1A) (Wu *et al*., 1998; Vale, 2003; Desnos *et al*., 2007b; Ross *et al*., 2008a; Hammer and Sellers, 2012; Heaslip *et al*., 2014; Fourriere *et al*., 2020; Pimm and Henty-Ridilla, 2021). In general, these vesicles undergo initial long-range transport from the Golgi by kinesin-1 motors on microtubules (MTs), with handoff to myosin Va motors for final, short-range transport on cortical actin filaments, where vesicles are delivered to the plasma membrane for secretion (Desnos *et al*., 2007a; Hammer and Sellers, 2012). Vesicular handoff between the MT and actin cytoskeletal systems occurs in cellular domains where kinesin-1 and myosin Va motors simultaneously engage their respective cytoskeletal tracks and compete mechanically to determine outcomes at actin-MT intersections. Thus, secretory vesicle transport involves actin-MT “crosstalk” that may be modulated to meet the cell’s physiological demands by ensuring vesicle delivery with spatial and temporal fidelity (Wu *et al*., 1998; Dogterom and Koenderink, 2019).

**Figure 1.**
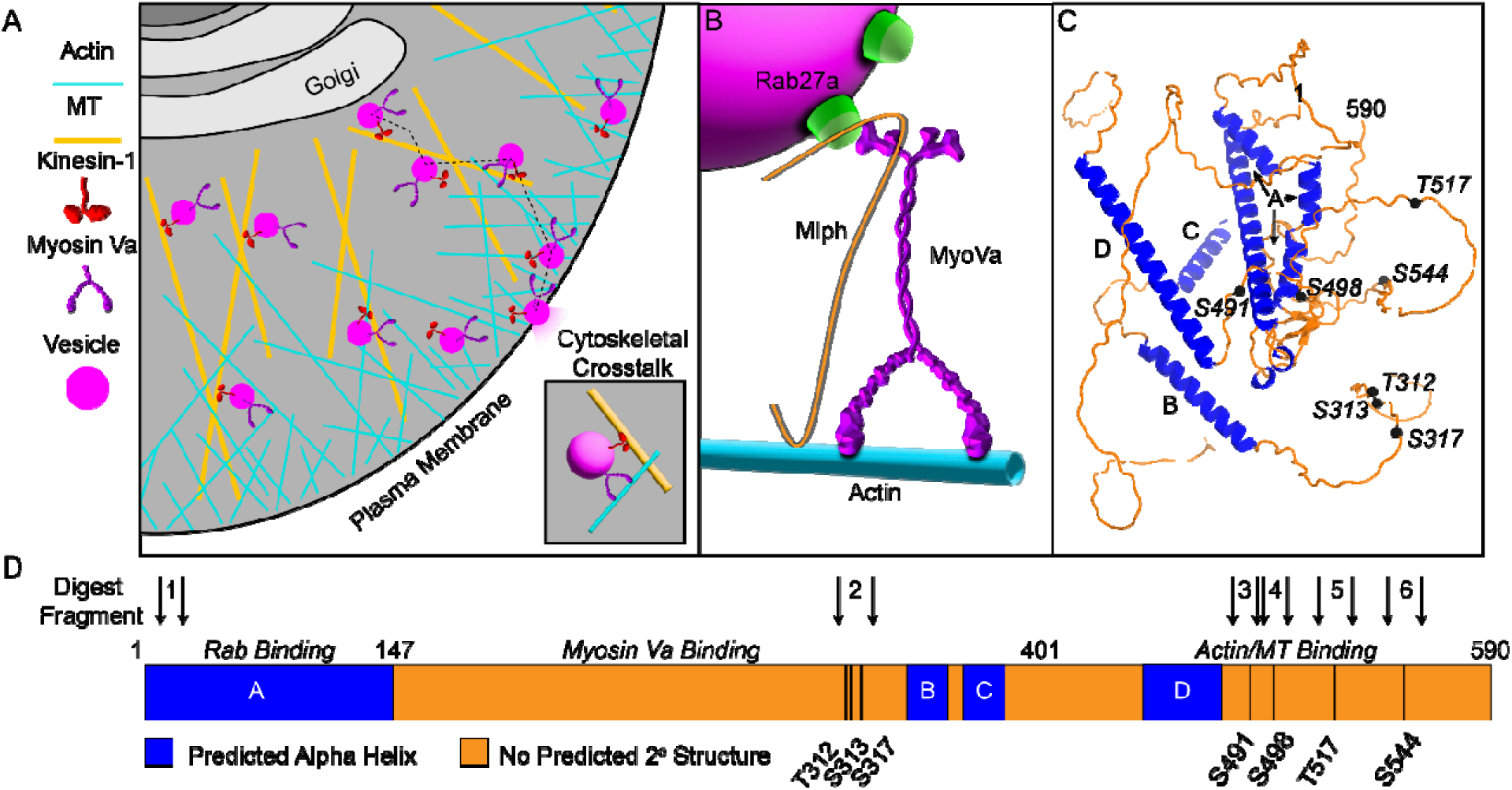
Structure and Function of the Myosin Va Adapter, Melanophilin. A) Cartoon depicting the cellular pathway along microtubules and actin filaments secretory vesicles take from the Golgi to the plasma membrane encountering numerous actin-microtubule intersections along the way. Inset: Illustration of cytoskeletal crosstalk which occurs when myosin Va and kinesin-1 motors attached to the same vesicle interact with their respective tracks. B) Illustration of how melanophilin (Mlph, orange) attaches Myosin Va (purple) to a vesicle cargo (magenta) via its interaction with Rab27a (green). Melanophilin also has a well-documented ability to bind to actin filaments (cyan). C) AlphaFold melanophilin structure prediction. Predicted alpha helical domains are shown in blue and labeled as in (D). Residues where no clear structure is predicted are shown in orange. Literature-supported potential phosphorylation sites are shown as black spheres and labeled as in (D). D) Primary melanophilin structure, with binding partner domains identified. Predicted alpha helical domains shown in blue and domains with no clear structural prediction shown in orange. Black vertical lines in the sequence indicate literature-supported potential phosphorylation sites with amino acid number below. Arrowheads indicate melanophilin digest peptides identified as phosphorylated by mass spectrometry with peptides numbered as in Table 1.

**Table 1:** Melanophilin phosphorylation varies with Protein Kinase A and Lambda Phosphatase treatment. Sf9-expressed melanophilin samples were either ‘Untreated’ or treated with Protein Kinase A (PKA) or lambda phosphatase (Dephos). Various melanophilin phosphopeptides were detected by mass spectrometry. For identification - sites in **bold** were unequivocally identified based on the y- and b-ion MS/MS spectra. Sites in *italics* are potential phosphorylation sites. *<u>Underlined</u>* sites are potential phosphorylation sites when the peptide contains a single phosphorylation. For quantification - ‘+’ indicates that a phosphopeptide was detected in the fragment. For the fragment H301-R322, a mixture of singly and doubly phosphorylated species were detected, so the ‘++’ indicates the dominant species. ND indicates that no phosphopeptide was detected.

| *Melanophilin Tryptic Peptides* | *Melanophilin Treatment* |  |  |
| --- | --- | --- | --- |
|  | **Untreated** | **PKA** | **Dephos** |
| 1) LDLSTLTDEEAEHVWAVVQR (L5-R24) | + | + | ND |
| 2) HLPSQYLADVD<u>T</u>SDEDSIQGPR (H301-R322) | Single: ++ | Single: + | ND |
|  | Double: + | Double: ++ | ND |
| 3) AAGLTVKPSGKPR (A483-R495) | ND | ND | ND |
| 4) RKSGIPIFLPR (R496-R506) | ND | + | ND |
| 5) IPKTPPADPDDQAK (I514-K527) | + | + | ND |
| 6) KY**SP**SSQGVDSGSFDR (K542-R557) | + | + | ND |

The simplest mode of biasing competition between kinesin-1 and myosin Va at actin-MT intersections is through tuning motor activity, with kinesin-1 and myosin Va sharing a fundamental regulatory paradigm where motors adopt a folded, autoinhibited conformation (Cai *et al*., 2007; Trybus, 2008; Verhey and Hammond, 2009; Tan *et al*., 2023; Carrington *et al*., 2024). These motors are activated when they bind to adapter proteins, which in some cases link the motor to membrane-bound Rab family GTPases, forming a tripartite complex on the vesicle surface (Hammer and Wu, 2002; Wu *et al*., 2002; Seabra and Coudrier, 2004). One such adapter is melanophilin, which links the myosin Va motor to Rab27a on melanosomes (Fig. 1B) (Wu *et al*., 1998, 2002), pigmented organelles that are trafficked by both kinesin-1 and myosin Va motors (Bao *et al*., 2025). Interestingly, beyond physically linking the myosin Va motor to the melanosome surface, melanophilin binds electrostatically to actin filaments *in vitro* via positively-charged amino acids in its C-terminus (Sckolnick *et al*., 2013; Oberhofer *et al*., 2017), thereby serving as a tether to enhance myosin Va transport distances. Melanophilin also binds the MT tip-tracking protein EB-1 (Wu *et al*., 2005), as well as MTs directly (Oberhofer *et al*., 2017, 2020), providing a potential mechanism for crosstalk between actin filaments and MTs. With multiple potential Protein Kinase A (PKA) sites in the C-terminus of melanophilin that may be phosphorylated *in vivo* (Fig. 1C,D) (Oberhofer *et al*., 2017), the potential for phosphorylation-dependent regulation of melanophilin’s binding to actin or MTs exists (Sheets *et al*., 2007; Oberhofer *et al*., 2017, 2020). In fact, Okten and coworkers have shown *in vitro* that phosphorylated melanophilin prefers actin, while dephosphorylated melanophilin preferentially binds MTs (Oberhofer *et al*., 2017, 2020), and that this preference biases track selection by single Rab27a/melanophilin/myosin Va complexes. However, melanosomes are transported by both myosin Va and kinesin-1 motors (Bao *et al*., 2025). Therefore, if melanophilin can potentially bias cargo track selection at an actin-MT intersection (Oberhofer *et al*., 2017, 2020), then this capacity would be best defined in the context of a physiologically relevant cargo transported by both myosin Va and kinesin-1 motors. Here, we investigate whether melanophilin-mediated cytoskeletal crosstalk biases the directional outcomes of liposomes being transported by both myosin Va and kinesin-1 motors at actin-MT intersections, and whether this bias is regulated by melanophilin phosphorylation.

Using an *in vitro* model of melanosome transport, we show that melanophilin can act as a phosphorylation-dependent switch that biases myosin Va/kinesin-1 liposomes at actin–MT intersections. In the absence of melanophilin, liposomes tend to remain on the track they entered the intersection on. Melanophilin overrides this bias in a phosphorylation-dependent manner and mirrors the track-binding preferences of individual melanophilin molecules. Our results suggest distinct mechanisms for directing transport, with melanophilin acting as an actin “anchor” that reinforces myosin Va engagement in contrast to a MT “tether” that promotes kinesin-1 engagement. These distinct melanophilin properties bias the tug-of-war between these motors at an actin-MT intersection, as might be the case *in vivo*. Therefore, rather than simply coupling motors to cargo, melanophilin can tune a cargo’s directional outcome at cytoskeletal intersections, providing a potential mechanism for redirecting cargo in response to cellular signals.

## Results

### Mass spectrometry analysis of Sf9-expressed melanophilin phosphorylation state in vitro

In melanophores, pigmented melanosomes either aggregate near the nucleus or disperse throughout the cell in response to PKA activation (Rogers *et al*., 1997; Sköld *et al*., 2002; Nascimento *et al*., 2003; Sheets *et al*., 2007), a redistribution that involves both actin- and MT-based motors (Rogers *et al*., 1997; Wu *et al*., 1998; Sköld *et al*., 2002; Sheets *et al*., 2007). Because melanosome-associated melanophilin can bind to both cytoskeletal tracks with a preference that depends on its phosphorylation state, we and others hypothesized that melanophilin phosphorylation could bias cargo routing at actin-MT intersections (Oberhofer *et al*., 2017, 2020) and thereby contribute to melanosome positioning in the cell. To test this possibility *in vitro*, we purified recombinant melanophilin expressed in Sf9 cells (Sckolnick *et al*., 2013) and altered its phosphorylation state by PKA or lambda phosphatase treatment in vitro (see Methods). Following tryptic digestion, at least 6 phosphorylated residues were identified by liquid chromatography mass spectrometry (LCMS), and the non-phosphorylated version of a peptide that contained a previously identified phosphorylation site (Table 1; Fig. 1C, D).

Untreated melanophilin showed basal levels of phosphorylation at 5 sites within 4 distinct peptides. Specifically, the most N-terminal peptide, L5-R24, which does not contain any reported phosphorylation sites, did show phosphorylation. Fragment H301-R322 contained 3 literature-supported phosphorylation sites (T312, S313, and S317) (Lundby *et al*., 2012; Carrier *et al*., 2016). These sites showed high occupancy in the singly phosphorylated state, based on a semi-quantitative mass balance approach (see Methods). Furthermore, peptide I514-K527, which contained a single literature referenced site at T517 (Zanivan *et al*., 2008), showed phosphorylation, as did peptide K542-R557, with its single literature referenced site, S544 (Zanivan *et al*., 2008; Oberhofer *et al*., 2017).

Following PKA treatment there was an apparent increase in melanophilin’s net phosphorylation, with phosphorylation sites identified in 5 peptides. The most dramatic increases occurred in peptide H301-R322, which shifted to mostly doubly phosphorylated species, while peptide R496-R506, with its well documented S498 phosphorylation site (Zanivan *et al*., 2008; Oberhofer *et al*., 2017) now clearly phosphorylated following PKA treatment (Table 1). Even though S491 in fragment A483-R495 is a predicted PKA site (Oberhofer *et al*., 2017), there was no detectable phosphorylation before or after PKA treatment, as previously reported (Oberhofer *et al*., 2017).

To generate a completely dephosphorylated melanophilin, we took advantage of an S491A/S498A double mutant that was reported to reduce PKA-dependent phosphorylation by ∼75% (Oberhofer *et al*., 2017). Lambda phosphatase treatment of this mutant melanophilin yielded protein with no detectable phosphorylation sites. Therefore, we effectively generated a gradient of melanophilin phosphorylation with PKA-treated being the highest, untreated being intermediate, and lambda phosphatase treated showing no detectable phosphorylation.

### Melanophilin phosphorylation state biases the track on which kinesin-1/myosin Va transported liposomes exit an actin-MT intersection

To understand the role of melanophilin’s phosphorylation state in biasing track selection at actin-MT intersections, we developed an *in vitro* model of cargo transport using fluid-like, 350-nm diameter liposomes decorated with on average 5 constitutively active kinesin-1 (K543), 5 constitutively active myosin Va (MyoVa HMM), and 5 melanophilin. The number of molecules on each liposome was estimated from a standard binding curve derived via a photobleaching-based counting method as described previously (Nayak and Rutenberg, 2011; Nelson *et al*., 2014; Bensel *et al*., 2024, 2025) (Fig. S1). These liposomes were then challenged with actin-MT intersections, which were assembled on a glass surface using a sequential flow technique such that the top and bottom tracks are known (see Methods) (Fig. 2A, E). We restricted our analysis to include only liposomes that entered the intersection starting on the bottom track and therefore were obstructed by the crossing track (Ross *et al*., 2008b; Schroeder *et al*., 2012; Lombardo *et al*., 2017; Deeb *et al*., 2019; Bensel *et al*., 2024). Under this condition, MyoVa HMM and K543 motors on the liposome could simultaneously engage the actin and MT in the intersection, respectively, and mechanically compete in a tug-of-war to decide the intersection outcome. For each actin-MT intersection encounter, we determined whether the liposome entered the intersection on actin or a MT, the track the liposome exited on, whether the liposome paused in the intersection, and the pause lifetime (Movies S1-S4).

**Figure 2.**
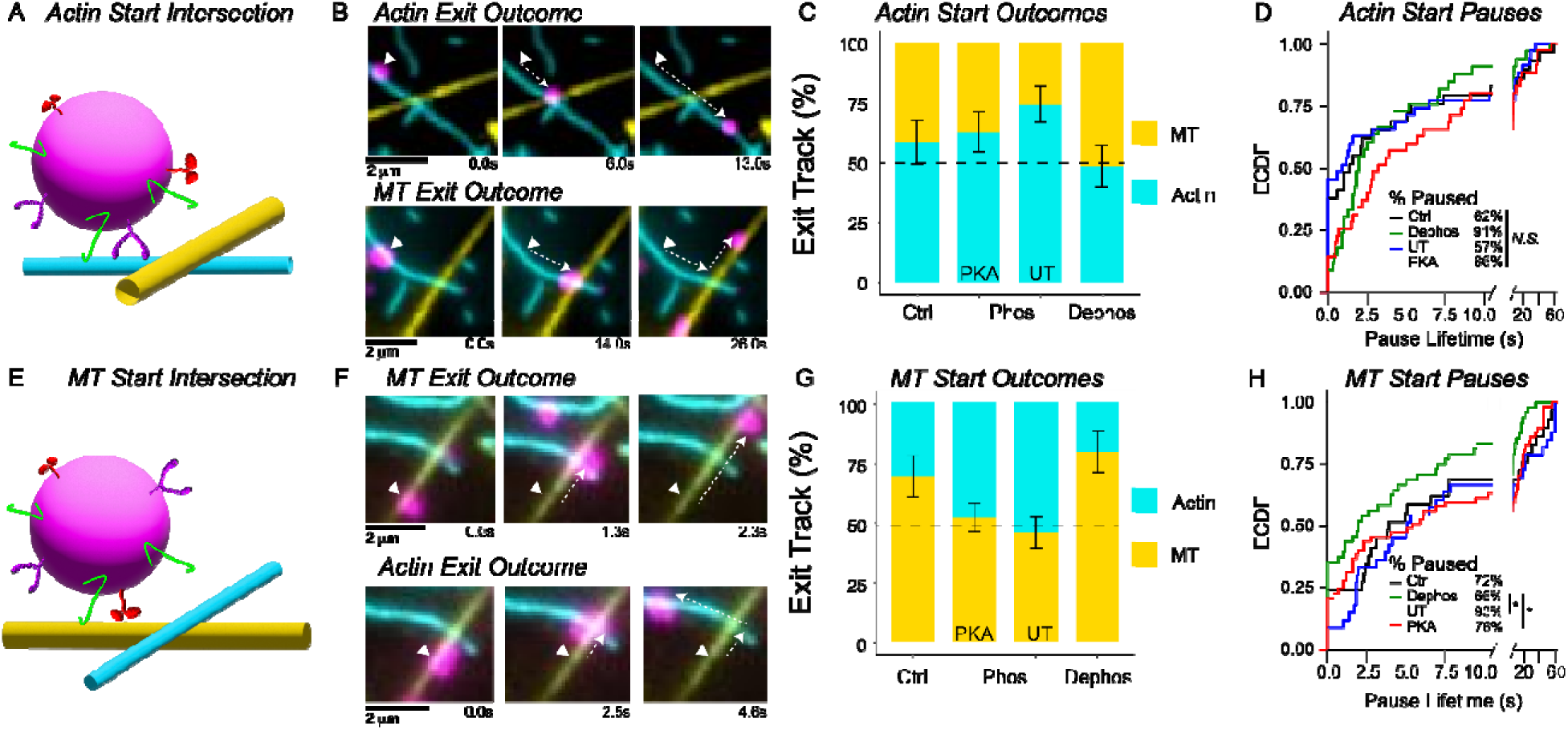
Melanophilin Phosphorylation State Biases Mixed-Motor Liposome Track Choice in Actin-MT Intersections. A) Schematic illustration of a myosin Va HMM (purple) and K543 (red) decorated liposome (magenta) with melanophilin (green) on the liposome surface encountering an actin (cyan)-MT (yellow) intersection with the liposome starting on actin. B) Example micrographs showing a mixed motor liposome starting on actin and then exiting an actin-MT intersection either on the actin filament it entered the intersection on (upper images) or on the MT (lower images). The liposome trajectory (dashed white arrow) and its starting point (white triangle) are identified. C) Bar graph depicting the percentage of actin-MT intersection exits on actin (cyan) or MTs (yellow) when the liposome enters the intersection on actin (as in A). The various conditions are mixed motor liposomes with no melanophilin present as a control (ctrl), mixed motor liposomes with PKA-phosphorylated melanophilin (PKA), mixed motor liposomes with untreated (UT) basally phosphorylated melanophilin, or mixed motor liposomes with lambda phosphatase-treated melanophilin (Dephos). Dashed horizontal line highlights 50% point. Error bars indicated standard deviation of bootstrapping analysis (2000 bootstrap replicates). N = 30-60 events per condition from 3 independent trials each. D) Empirical Cumulative Distribution Function (ECDF) plot of pause lifetimes observed in actin start actin-MT intersections for the various conditions as abbreviated in C and color-coded in the inset, where the percentage of intersection encounters for a given condition with a detectable pause (3+ consecutive frames of no motion in the intersection) is shown. Statistical analysis was performed by a Kruskal-Wallis test in R, N.S. = p>0.05. E) Schematic illustration of mixed-motor intersection assay where the liposome enters the intersection on the MT, which is now the bottom track. F) Example micrographs showing a mixed motor liposome starting on MT and then exiting an actin-MT intersection on the MT it entered the intersection on (upper images) or on the actin filament (lower images). The liposome trajectory (dashed white arrow) and its starting point (white triangle) are identified. G) Bar graph depicting the percentage of actin-MT intersection exits on actin (cyan) or MTs (yellow) when the liposome enters the intersection on MT (as in E). The various conditions and bar chart features are as described in C. H) Empirical Cumulative Distribution Function (ECDF) plot of pause lifetimes observed in MT start actin-MT intersections for the various conditions as abbreviated in C and color-coded in the inset, where the percentage of intersection encounters for a given condition with a detectable pause (3+ consecutive frames of no motion in the intersection) is shown. Statistical analysis was performed using a Kruskal-Wallis test with Dunn’s post hoc test in R, *, p<0.05, all other pairwise comparisons have p>0.05.

Control liposomes without melanophilin (Movies S2-S4), regardless of whether they entered the actin-MT intersection on the actin filament or MT, show a bias towards exiting the intersection on the same track they entered on (Fig. 2C, G; Ctrl). In both cases, the majority of liposomes (>60%) paused in the intersection (median lifetime: actin, 1.4s; MT, 3.9s; Fig. 2D, H; Ctrl), suggesting that a tug-of-war between actin-engaged myosin Va and MT-engaged kinesin occurred in the intersection (Ross *et al*., 2008b; Schroeder *et al*., 2012; Osunbayo *et al*., 2015; Lombardo *et al*., 2017; Bensel *et al*., 2024).

To determine if the presence of melanophilin on the liposome surface and its phosphorylation state dictate on which track the liposome exited the actin-MT intersection, we used motor-decorated liposomes with an average of 5 melanophilin molecules that were either PKA-phosphorylated, untreated, or completely dephosphorylated with lambda phosphatase (see Methods). Because MyoVa HMM is constitutively active and lacks its melanophilin binding domain, the presence of melanophilin on the liposome surface gives specific insight into how melanophilin’s track binding directly impacts liposome transport. For liposomes entering the intersection on actin (Movies S1, S2), whether surface-bound melanophilin was PKA-phosphorylated or basally phosphorylated (i.e., untreated) (Table 1), there remained a bias towards exiting the intersection on the actin filament as in the absence of melanophilin (Fig. 2C; Phos). However, this bias was lost with dephosphorylated melanophilin, so exiting on either track was similarly likely (Fig. 2C; Dephos). When entering the intersection on a MT (Movies S3, S4), liposomes with either PKA-phosphorylated or untreated melanophilin also showed no track preference when exiting the intersection (Fig. 2G; Phos). In contrast, the presence of dephosphorylated melanophilin further enhances the bias to exit the actin-MT intersection on MTs (Fig. 2G; Dephos).

As for pause lifetimes in the intersection, liposomes starting on actin had no significant differences as a function of melanophilin’s phosphorylation state (Fig. 2D). Whereas for liposomes entering the intersection on a MT, liposomes with dephosphorylated melanophilin showed shorter pause lifetimes (median = 2.0s) compared to those with PKA-phosphorylated (median = 6.0s) or untreated melanophilin (median = 5.1s) (Fig. 2H). Taken together, these data suggest that the presence of melanophilin on a liposome and its phosphorylation state modulates the preferred track on which the liposome exits the actin-MT intersection (i.e., phosphorylated melanophilin biases liposomes towards actin while dephosphorylated melanophilin biases liposomes towards MTs) with the duration of pausing in actin-MT intersections indicative of a tug-of-war that is dependent on melanophilin’s phosphorylation state.

### Melanophilin’s phosphorylation state biases track binding preference

To determine melanophilin’s track preference and whether it changes with melanophilin’s phosphorylation state, we measured the *in vitro* binding of Qdot-labeled melanophilin molecules to both actin filaments and MTs bound to the same glass surface (Fig. 3A). PKA-phosphorylated melanophilin preferentially bound to actin filaments versus MTs (0.42 bound/μm actin vs. 0.18 bound/μm MT; Fig. 3B, F, Movie S5), whereas untreated melanophilin showed no clear preference (0.45 bound/μm actin vs. 0.43 bound/μm MT; Fig. 3C, F, Movie S6). In contrast, dephosphorylated melanophilin preferentially bound to MTs (0.34 bound/μm actin vs. 0.63 bound/μm MT; Fig. 3D, F, Movie S7). These results suggest that melanophilin’s phosphorylation state predominantly alters MT binding which gradually decreases with increasing phosphorylation, while actin binding was unaffected (Fig. 3F) as reported previously (Oberhofer *et al*., 2017, 2020).

**Figure 3.).**
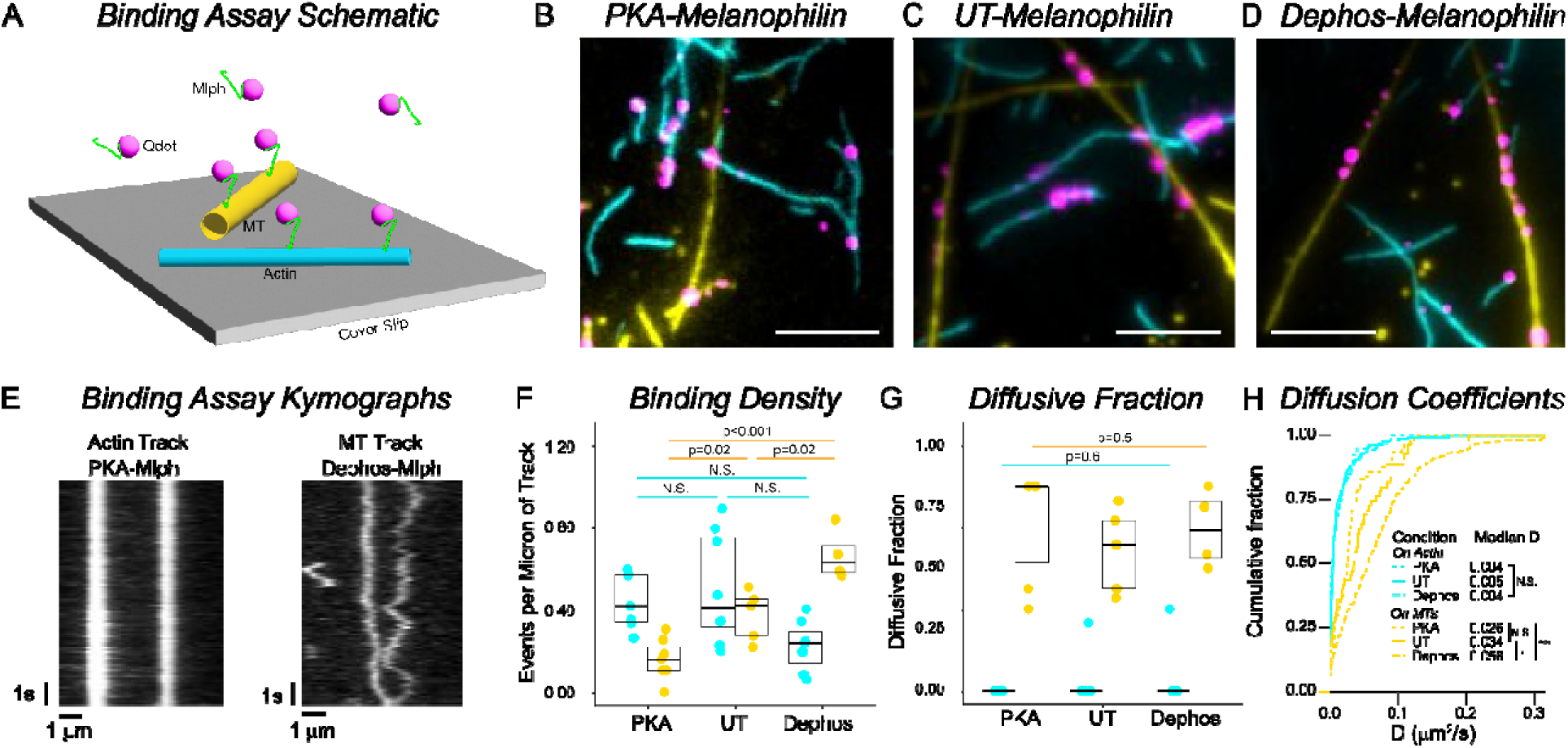
Melanophilin’s Phosphorylation State Modulates its Track Binding Preference In Vitro. A) Schematic illustration of *in vitro* melanophilin binding assay. Single melanophilin molecules (green) were labeled with fluorescent Qdots (magenta) and introduced into a flow cell with both actin filaments (cyan) and MTs (yellow) affixed to the cover slip (grey) surface, then imaged with TIRF microscopy. B) Example micrograph of binding assay performed with PKA-phosphorylated melanophilin. Qdots, actin, and MTs are colored as in A. Scale bar = 5 μm. C) Example micrograph of binding assay performed with untreated (UT) melanophilin color coded as in A. Scale bar = 5 μm. D) Example micrograph of binding assay performed with dephos-melanophilin after lambda phosphatase treatment color coded as in A. Scale bar = 5 μm. E) Kymograph of PKA-treated melanophilin interacting with actin (left) and dephosphorylated melanophilin interacting with MTs (right). Time is on the vertical axis and distance along the track is shown on the horizontal axis. F) Dot plot of melanophilin binding density on actin (cyan) and MTs (yellow) with overlaid box plot for the various conditions: PKA-phosphorylated (PKA), untreated (UT), and lambda phosphatase-treated (Dephos). Each dot represents a single kymograph where the binding density is defined as N(bound Qdots)/filament length (μm). Statistical analysis is a Kruskal-Wallis test using Dunn’s post-hoc test for pairwise comparisons. G) Dot plot of diffusive fraction of bound Qdot-labeled melanophilin on actin (cyan) and MTs (yellow) with overlaid box plot for condition as in F. Each dot represents a single kymograph where diffusive fraction is defined as N(diffusing Qdots)/N(total bound Qdots). Statistical analysis performed using a Kruskal-Wallis test. H) Cumulative distribution plots depicting measured diffusion coefficients (D) for individual Qdot-labeled melanophilin tracked along either actin filaments (cyan) or MTs (yellow). Each condition is signified by line type as defined in the inset, where median D values per condition and statistical analysis are shown. Statistics: ***, p<0.001, *, p<0.05, with D value distributions compared by the Kolmogorov-Smirnov test with Holm’s adjustment for multiple comparisons.

Once bound to either track, melanophilin molecules generally remained bound for the length of the recorded video (i.e., 10s). Nearly all melanophilin bound statically to actin regardless of melanophilin’s phosphorylation state (Fig. 3E, G) with a diffusion constant ≤0.005 μm^2^/s (Fig. 3H), equivalent to our tracking error defined as the predicted length of the melanophilin molecule plus our localization error (Pouget *et al*., 2004; Nelson *et al*., 2006; Michalek *et al*., 2015). Whereas nearly all MT-bound melanophilin diffused along the MT lattice (Fig. 3E, G; Movies S5-S7), indicative of a fundamental difference in track interaction. The diffusion constants on MTs for PKA-phosphorylated (0.026 μm^2^/s) and untreated melanophilin (0.034 μm^2^/s) were similar but slower than that for dephosphorylated melanophilin (0.056 μm^2^/s) (Fig. 3H).

### Melanophilin phosphorylation modulates liposome transport on single tracks

As described above (Fig. 2), when a liposome transported by a mixed population of kinesin-1 and myosin Va motors encounters an actin-MT intersection, the track it exits on can be biased by the presence and phosphorylation state of melanophilin on the liposome surface. This tuning of intersection outcomes mechanistically could relate to both melanophilin’s track affinity (see above, Fig. 3) and how such track binding impacts the apparent transport capabilities of the motors themselves (Ross *et al*., 2008a; Bensel *et al*., 2025). Therefore, we first characterized multimotor liposome transport (Fig. 4A and E) (i.e. run length, velocity, frequency of reaching the track end) by ∼5 MyoVa HMM on single actin filaments or by ∼5 K543 on single MTs in comparison to a single Qdot-labeled motor (Fig. 4). These multimotor liposome transport data were then compared to transport of similar liposomes but with ∼5 melanophilin on the liposome surface that were PKA-phosphorylated, untreated, or dephosphorylated (Fig. 4).

**Figure 4.**
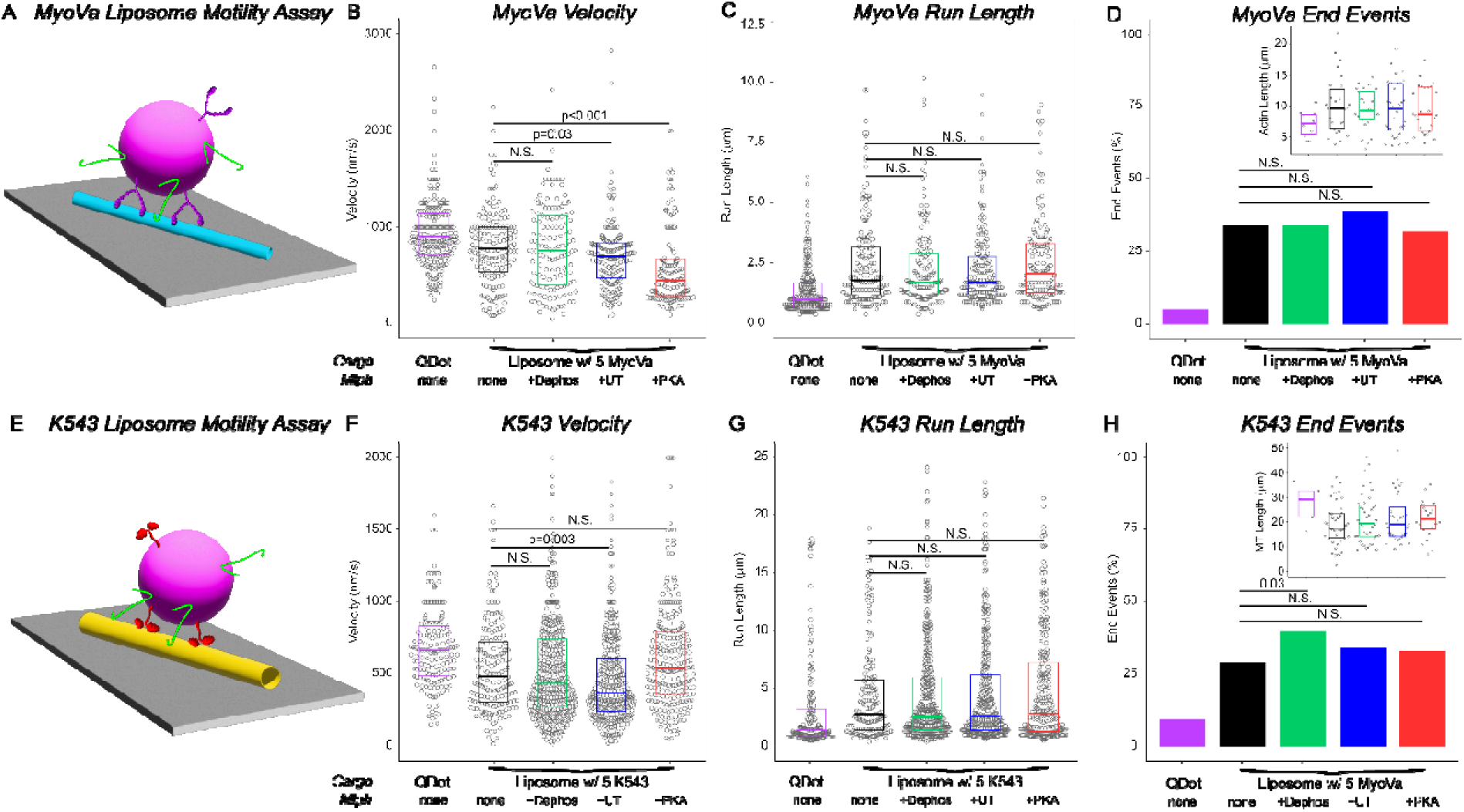
Melanophilin Phosphorylation State Modulates Liposome Transport Along Single Tracks In Vitro. A) Illustration of liposome motility assay on an actin filament. Liposome (magenta) decorated with ∼5 myosin Va HMM (purple) and ∼5 melanophilin (green) are introduced to a flow cell with actin (cyan) affixed to a cover slip (grey) and are imaged with TIRF microscopy. B) Dot plot of myosin Va HMM velocity with overlaid box plot. Each dot represents a single event. Conditions are a Qdot-labeled single myoVa HMM molecule (Qdot, purple), myoVa HMM liposomes with no melanophilin as a control (none, black), myoVa HMM liposomes with lambda phosphatase treated melanophilin (+Dephos, green), myoVa HMM liposomes with untreated melanophilin (+UT, blue), and myoVa HMM liposomes with PKA-phosphorylated melanophilin (+PKA, red). Statistical analysis was a Kruskal-Wallis with Dunn’s post-hoc test for pairwise comparisons. C) Dot plot of myosin Va HMM run lengths with overlaid box where each dot represents a single event. Conditions and statistics as in B. D) Bar graph of the frequency at which myosin Va HMM-transported cargos reach the actin filament end without detaching (End Events). Bars colors match that in B and C. Statistical analysis used a pairwise Proportion Test in R with the Benjamini-Hochberg correction for multiple comparisons. Inset: Dot plot of actin filament lengths from which the data were collected for each condition. Colors match those in the main panel. E) Illustration of liposome motility assay on a MT. Liposomes (magenta) decorated with ∼5 K543 (red) and ∼5 melanophilin (green) are introduced to a flow cell with MTs (yellow) affixed to a cover slip (grey) and are imaged with TIRF microscopy. F) Dot plot of K543 velocity with overlaid box plot, where each dot represents a single event. The various conditions and statistics are as described in B. G) Dot plot of K543 run lengths with overlaid box plot where each dot represents a single event. The various conditions and statistics are as described in B. G) Bar graph of the frequency at which K543-transported cargos reach the end of the MT without detaching (End Events). Bars colors match that in F and G. Statistical analysis used a pairwise Proportion Test in R with the Benjamini-Hochberg correction for multiple comparisons. Inset: Dot plot of MT lengths from which the data were collected for each condition. Colors match those used in the main panel.

Liposomes with ∼5 MyoVa HMM have significantly longer run lengths and slower velocities than single Qdot-labeled MyoVa HMM (Fig. 4B, C), as expected for multimotor transport (Nelson *et al*., 2014). Interestingly, the addition of ∼5 PKA-phosphorylated, untreated, or dephosphorylated melanophilin to these liposomes did not significantly change the median run length (Fig. 4C). However, with a high frequency of liposomes reaching the end of the actin filament (Fig. 4D), the true run length may be underestimated compared to what would be possible on an infinitely long actin filament (Mayr *et al*., 2011; McKenney *et al*., 2014; Nelson *et al*., 2014; Soppina *et al*., 2014; Bensel *et al*., 2024). Therefore, the frequency of liposomes reaching the end of the track can reveal otherwise undetectable changes in true run length. With there being no difference in the frequency at which the liposome reached the end of the actin filament under all experimental conditions (Fig. 4D), and distributions of actin filament lengths not being significantly different (Fig. 4D inset), then the lack of an effect of melanophilin and its phosphorylation state on liposome run length is assumed real. Interestingly, liposome velocity (Fig. 4B) was reduced by PKA-phosphorylated melanophilin compared to the no melanophilin control (median±SEM, 450±40 vs. 780±30 nm/s, respectively) and to a lesser extent by untreated melanophilin (median±SEM, 700±30 vs 780±30 nm/s, respectively), while dephosphorylated melanophilin showed no effect.

Liposomes transported by ∼5 K543 motors show significantly longer run lengths and slower velocities than single Qdot-labeled K543 motors (Fig. 4F, G), once again as expected for multimotor transport (Kunwar and Mogilner, 2010; Derr *et al*., 2012; Furuta *et al*., 2013; Grover *et al*., 2016; Feng *et al*., 2018; Tjioe *et al*., 2019; Sarpangala and Gopinathan, 2022; Bensel *et al*., 2024). While there is no statistically significant change in the multimotor liposome median run length with PKA-phosphorylated, untreated, or dephosphorylated melanophilin (Fig. 4G), there in fact may be differences if the melanophilin phosphorylation state alters the frequency of liposomes reaching the MT end. With the MT length distributions for each of the experimental conditions not being different (Fig. 4H, inset), the fact that dephosphorylated melanophilin significantly increases the probability of reaching the MT end (41% vs. 29% control), which was not the case for PKA-phosphorylated or untreated melanophilin (Fig. 4H), suggests that dephosphorylated melanophilin does increase the liposome run length. As for liposome velocities, untreated melanophilin slightly reduced liposome velocity on MTs while PKA-phosphorylated and dephosphorylated melanophilin did not, compared to control (Fig. 4F).

## Discussion

Myosin Va and kinesin-1 motors cotransport secretory vesicles from the cell center towards the plasma membrane (Varadi *et al*., 2002, 2005; Desnos *et al*., 2007b; Ross *et al*., 2008a; Hammer and Sellers, 2012). In doing so, they maneuver their shared cargo through numerous actin-MT intersections (Fig. 1A) (Heaslip *et al*., 2014; Dogterom and Koenderink, 2019; Pimm and Henty-Ridilla, 2021). Given the physiological need for cargo delivery to be spatially and temporally correct, mechanisms must exist to bias track selection at these intersections. Here we show using a simplified *in vitro* cargo transport model system that melanophilin, a myosin Va adapter protein, when on the surface of mixed myosin Va and kinesin-1 transported liposomes, provides a means of biasing track selection at actin-MT intersections that is dependent on melanophilin’s phosphorylation state (Fig. 2C, G), as predicted previously based on single myosin Va motor complex interactions with actin and MT tracks (Oberhofer *et al*., 2017, 2020). Our liposome model system with its attached myosin Va and kinesin-1 motors allows melanophilin tip the scales of the tug-of-war that must exist *in vivo* at actin-MT intersections. Specifically, we observed that liposomes with phosphorylated melanophilin on their surface are biased toward exiting intersections on actin, whereas dephosphorylated melanophilin biases these exits to be on MTs. Interestingly, without melanophilin (Fig. 2C, G: Ctrl), liposomes prefer to remain on their track of entry, even though the intersecting track is a large obstacle that the transporting motors cannot step over. In this case, motors that are not actively transporting the liposome (i.e., MyoVa HMM or K543) are diffusing on the liposome surface and therefore are free to bind to the segment of the same track beyond the intersection before motors of the opposite type can fully engage the intersecting track. Such a track binding advantage is a potential mechanism that biases the liposome to exit the intersection on its track of entry.

Although we show that melanophilin on the liposome surface can bias track selection at actin-MT intersections, other mechanisms exist that modulate cargo transport which involve key elements of cargo transport systems. For example, track preference can be modulated by altering a motor’s track affinity through track-binding proteins such as MT-associated proteins (MAPs) (Vershinin *et al*., 2007; Lipka *et al*., 2016; Stern *et al*., 2017; Chaudhary *et al*., 2018; Bodakuntla *et al*., 2019; Pan *et al*., 2019; Monroy *et al*., 2020), tropomyosin isoforms on actin filaments (Gunning *et al*., 2008; Sckolnick *et al*., 2016), and septins (Spiliotis and Nakos, 2021), which can bind to both MTs and actin; a prime example of cytoskeletal crosstalk. Furthermore, if myosin Va and kinesin-1 motors can transition between a folded, autoinhibited state and an extended, active state while attached to cargo, this dynamic could then alter the number of available motors, biasing transport outcomes. For example, we previously showed that nearly full-length kinesin-1 motors on a liposome surface exist in a dynamic equilibrium between active and autoinhibited states which effectively reduced the number of motors available to engage the MT, impacting directional outcomes at MT-MT intersections (Bensel *et al*., 2025).

### Melanophilin phosphorylation impacts track binding affinity and molecular motor transport properties

Using a combination of single-molecule melanophilin track-binding assays and multimotor liposome transport assays with melanophilin present on the liposome surface (Figs. 3, 4), we determined melanophilin’s phosphorylation-dependent preference for actin versus MT binding and the effective strength of its association with each track once bound. Based on these assays, the extent of melanophilin’s actin binding was phosphorylation independent (Fig. 3F), in agreement with previous findings (Oberhofer *et al*., 2017). However, once actin-bound, phosphorylated melanophilin (both PKA-treated and untreated) behaved effectively as an “anchor” that resisted liposome transport by myosin Va, as evidenced by the slowing of transport velocity (Fig. 4B). Because myosin Va liposome transport was slower in the presence of PKA-treated melanophilin than basally phosphorylated melanophilin (Fig. 4B), the strength of melanophilin’s anchoring effect appears to scale with phosphorylation levels. We previously reported slowing of a single myosin Va linked to a Qdot-labeled melanophilin (Sckolnick *et al*., 2013) suggesting that melanophilin’s interaction with actin is sufficiently strong to create a resistive load against which the motor must operate. We also showed that, as the myosin Va stepped along actin, melanophilin’s C-terminus moved in discrete ∼108nm steps as it lagged behind the myosin Va (Sckolnick *et al*., 2013). This suggests that melanophilin binds strongly to discrete sites on actin, and that melanophilin is quite extensible, which is consistent with the Alphafold predicted structure (Fig. 1C). It is notable that slowing of myosin Va by melanophilin we report here and previously (Sckolnick *et al*., 2013), was not observed by Okten and coworkers (Oberhofer *et al*., 2017) using a tripartite complex of Rab27a, melanophilin, and myosin Va. One explanation is that the tripartite complex may have a constituent stoichiometry different than the assumed 1:1:1 making comparisons difficult.

Unlike its interaction with actin, melanophilin binding to MTs was strongly phosphorylation dependent, increasing progressively with decreasing phosphorylation (Fig. 3F). With dephosphorylation effectively making melanophilin more positively charged, the increased propensity of dephosphorylated melanophilin to bind the MT surface with its numerous negatively-charged tubulin E-hooks is expected (Minoura *et al*., 2010). Curiously, the diffusion constant of melanophilin shows the opposite relationship to that expected for a simple positively-charged protein or object on a MT (Minoura *et al*., 2010), suggesting that melanophilin’s diffusive interaction with MTs is far more complex. The clearest effect of melanophilin, a myosin Va adapter, on kinesin-1-driven liposome transport was that dephosphorylated melanophilin significantly increased the frequency with which kinesin-1-transported liposomes reached the ends of MTs (Fig. 4H). This suggests that dephosphorylated melanophilin functions as a tether, maintaining liposome engagement with the MT through the same C-terminal domain that binds actin (Oberhofer *et al*., 2017, 2020), while its diffusive interaction with the MT imposes little or no resistive drag. These distinct actin and MT binding preferences suggest that melanophilin’s interaction with these two tracks is fundamentally different, which is intriguing given that they are differentially regulated by the same phosphorylation sites (Oberhofer *et al*., 2017, 2020).

### Melanophilin’s phosphorylation state modulates crosstalk in actin-MT intersections by imparting advantages in the tug-of-war

Liposome cargos decorated with multiple molecules each of MyoVa HMM, K543, and melanophilin provide mechanistic insights into the tug-of-war between two motor systems at cytoskeletal intersections, as expected in cells (Verdeny-Vilanova *et al*., 2017), which would be impossible using single motor complexes. For example, liposome pausing at actin-MT intersections with pause lifetimes dependent on melanophilin’s phosphorylation state (Fig. 2D, H), suggest that melanophilin’s track binding properties contribute to biasing the eventual tug-of-war outcome and thus the track on which the liposome exits the intersection. Specifically, with phosphorylated melanophilin having a higher affinity for actin than MTs (Fig. 3F) and anchor-like behavior on actin (Fig. 4B) (Sckolnick *et al*., 2013), greater average force would be required of the MT-engaged K543 motors to win the tug-of-war, thus biasing the liposome to prefer actin exits. In contrast, with dephosphorylated melanophilin on the liposome, which preferentially associates with MTs (Fig. 3F) and promotes enhanced K543 MT engagement (see Results), the tug-of-war outcome is then biased so that liposomes prefer exiting the intersection on MTs. These findings from our simplified *in vitro* model system may contribute to understanding how cytoskeletal preference switching, based on the melanophilin phosphorylation state, dictates cellular localization of melanosomes between the MT-rich perinuclear region and the actin-rich cortex in response to PKA signaling (Rogers *et al*., 1997; Sheets *et al*., 2007; Bao *et al*., 2025).

### Cargo adapter proteins: an underappreciated mediator of cytoskeletal crosstalk

Here we describe how melanophilin, a myosin Va cargo adapter protein (Wu *et al*., 2002; Sheets *et al*., 2007), independent of its myosin Va activating role, is involved in cytoskeletal crosstalk to bias track preference for myosin Va/kinesin-1 transported liposomes at actin-MT intersections. Could this dual functionality be shared with other adapter proteins? Interestingly, constitutively active motors expressed in cells do not always accumulate in the same subcellular location as their cargos (Jenkins *et al*., 2012; Bentley and Banker, 2016), and cargos with different destinations are sometimes transported by the same motor (Fan and Lai, 2022; Wong and Weisman, 2021; Reck-Peterson *et al*., 2018). If constitutively active motors interpret track structure and its modifications in the same way as full length motors bound to cargo, then various vesicle-bound proteins may confer additional bias to track selection at the cargo level. In fact, granuphilin (Brozzi *et al*., 2012), rabphilin 3a (Brozzi *et al*., 2012), and MyRIP (Desnos *et al*., 2003), closely related myosin adapter proteins to melanophilin, are expressed primarily in neurons and endocrine cells where cargo transport and delivery is critical to normal function (Mizoguchi *et al*., 1994; Coppola *et al*., 2002; Fukuda and Kuroda, 2002). Could this family of cargo adapters provide cells with another tool for ensuring spatially and temporally specific cargo delivery? The work shown here with melanophilin paves the way for understanding the broader capacity of non-motor cargo-bound proteins as modulators of cytoskeletal crosstalk in a manner that is crucial for supporting biological function.

## Methods

### Expression, biotinylation, and purification of recombinant proteins

Full-length wild type *Mm* melanophilin (Sckolnick *et al*., 2013) (accession no. Q91V27.1) was used either untreated or phosphorylated with PKA (see below). A mutant *Mm* melanophilin with two site mutations, S491A and S498A (Oberhofer *et al*., 2017), was used for dephosphorylation by lambda phosphatase (see below). Both melanophilin constructs have N-terminal biotin and FLAG tags and were expressed using the baculovirus/Sf9 system and purified using FLAG affinity chromatography. Briefly, Sf9 cells are infected with recombinant baculovirus encoding melanophilin and grown in media supplemented with 0.2 mg/ml biotin for 72 hours, with the protein biotinylated during expression. Pelleted cells were resuspended in lysis buffer (10 mM imidazole, pH7.4, 0.3 M NaCl, 1 mM EGTA, 7% sucrose, 1 mM DTT, 0.5 mM AEBSF, 0.5 mM TLCK, 5 μg/ml leupeptin) and lysed by sonication and clarified by ultracentrifugation (200,000 x g for 40 min). The supernatant is then incubated with FLAG affinity resin for 1h at 4° C and then washed with Wash Buffer (10 mM, imidazole pH7.4, 0.3 M NaCl, 1 mM EGTA) and eluted with 0.1 mg/ml FLAG peptide in Wash Buffer. 1 mM DTT was added to pooled fractions of eluate prior to concentration over an Amicon 10K centricon and dialysis into Storage Buffer (10 mM imidazole, pH 7.4, 0.2 M NaCl, 1 mM EGTA, 2 mM DTT, 50% Glycerol). An additional construct of WT *Mm* melanophilin with a C-terminal YFP was used for photobleaching counting assays (see below).

K543 is a truncation of kinesin-1, *Mm* Kif5b (accession no. Q61768), that contains the N-terminal 543 amino acids of the native protein followed by C-terminal biotin and FLAG tags (Bensel *et al*., 2024). A second K543 construct with an N-terminal YFP was used for photobleaching counting assays. The K543 constructs were expressed and purified using the methods outlined above, with the addition of 2mM MgATP in the lysis buffer, and 100μM MgATP in the final storage buffer.

MyoVa HMM is a heavy meromyosin truncation of *Mm* myoVa (accession no. XP_006510890.1) truncated at amino acid 1098, followed by C-terminal biotin and FLAG tags (Hodges *et al*., 2007). An additional MyoVa HMM construct, with an N-terminal YFP was used for photobleaching motor counting assays. Both MyoVa HMM constructs were co-expressed with a calcium-insensitive calmodulin mutant (accession no. NP_001008160 with the mutations E32Q, E68Q, E105Q, and E141Q) and purified using the methods outlined above with the addition of 2mM MgATP in the lysis buffer.

### Treatment of melanophilin and characterization by liquid chromatography mass spectrometry (LCMS)

PKA-treated melanophilin was generated by phosphorylating 135 μg of WT melanophilin with 87,500U of cAMP-dependent Protein Kinase A (PKA) catalytic subunit (NEB P6000S) with 2 mM MgATP at 30° C for 1 hour, then incubation on ice overnight. Dephosphorylated melanophilin was generated by treating 100 μg mutant melanophilin (S491A/S498A) with 1000U of Lambda Phosphatase (NEB P0753S) at 37° C for 30 minutes, then incubation on ice overnight.

In preparation for LCMS, approximately 10 μl of melanophilin was solubilized in Rapigest surfactant, reduced with dithiothreitol, alkylated with iodoacetamide, and digested overnight with trypsin to generate peptides, as previously described (O’Leary *et al*., 2019). Following digestion, Rapigest was degraded using formic acid and trifluoroacetic acid, and peptides were collected for analysis (O’Leary *et al*., 2019). Peptides were separated by ultra-high-performance liquid chromatography using an XSelect HSS T3 1 x 150 mm x 3.5 µm column (Waters) and analyzed on a Q Exactive Hybrid Quadrupole-Orbitrap mass spectrometer (Thermo Fisher Scientific) operated in data-dependent MS mode, with the five most abundant precursor ions selected for fragmentation (Wood *et al*., 2022).

#### Data Analysis

Raw LCMS data files were processed using Thermo Proteome Discoverer (PD v2.2.0.388). Mass spectra were searched with Sequest HT against a custom mouse melanophillin protein sequence combined with the Mus musculus UniProt reference proteome (74,085 sequences; downloaded 02/09/2015). LCMS peak areas were generated using PD and normalized to the abundance of the top 3 ionizing peptides (AFIEVGQK, ELLSDTAHLNETHCAR, AAGLTVKPSGKPR) in Excel (Microsoft) to account for differences in column loading. The degree of phosphorylation was estimated from the change in the abundance of the non-phosphorylated variant of the peptide where available, using a mass-balance approach as previously described (Previs *et al*., 2008).

### Preparation and characterization of liposome cargos

Liposomes were prepared as described previously (Nelson *et al*., 2014; Lombardo *et al*., 2017; Bensel *et al*., 2024). Briefly, a lipid mixture (described in molar ratio) of 84 parts DOPC, 5 parts PEG-ylated DOPE, 5 parts cholesterol, 5 parts MBP:PE, and 1 part lipophilic fluorescent dye DiD was initially dried under N_2_ gas followed by 1 hour in vacuum. The lipid mixture was then rehydrated overnight in PBS (pH = 7.4) and extruded through a 1 μm pore size filter. 1 μM thiolated neutravidin (SH-NaV) was added to the extruded lipids and bound to the MBP:PE lipid overnight. Excess SH-NaV was washed away by ultracentrifugation (392,000 x g for 10 minutes) and liposomes were resuspended 3x in PBS. Finally, liposomes were extruded to their final size of ∼350nm through a 200-nm pore size filter. This preparation yields ∼250 μl of 10 nM liposomes.

To estimate the number of bound molecules per liposome, we use a photobleaching approach initially described by Nayak and Rutenberg (Nayak and Rutenberg, 2011). Our exact approach and data analysis are thoroughly described in Nelson et al., (Nelson *et al*., 2014) and Bensel et al (Bensel *et al*., 2024). Briefly, liposomes are prepared specifically for counting assays with the omission of the fluorescent DiD dye. These liposomes are mixed with varying stoichiometries of YFP-tagged K543, MyoVa HMM, or melanophilin ranging from 5 molecules per liposome to 20 molecules per liposome in Actin Buffer (25 mM Imidazole pH = 7.4, 25 mM KCl, 4 mM EGTA, 4 mM MgCl_2_). The resulting liposome-protein complexes are introduced into a flow cell and imaged in TIRF microscopy. When a liposome lands on the flow cell surface, there is a bright initial fluorescence followed by a gradual decay, creating a photobleaching transient (Fig. S1A). We collect a large number (>50) of transients for each condition to determine an average photobleaching transient. Then, for each individual transient, the intensity per fluor is mathematically related to the variance between that transient and the average transient, and the total number of fluors is given by the initial intensity divided by the intensity per fluor for any given transient (Fig. S1B). For K543 and MyoVa the number of fluors must be divided by 2 to account for the dimeric nature of these motors, but for melanophilin the number of fluors is equal to the number of molecules. From these measurements we create a standard curve of average number of bound molecules versus the incubation stoichiometry (melanophilin standard curve shown in Fig. S1C). Finally, we incubated liposomes with 5 MyoVa HMM, 5 K543, and 5 melanophilin per liposome and determined the average number of fluors per liposome, where we expect ∼25 YFP molecules per liposome and observed 25±3 (Fig. S1C).

### Actin-MT intersection assay

We used a TIRF microscopy approach similar to that described previously (Lombardo *et al*., 2017; Deeb *et al*., 2019; Bensel *et al*., 2024) but with modifications to utilize both actin and MTs and with surface passivation optimized for the cargos described here. Flow cells with two perpendicular channels were assembled using a silanized (Dixit and Ross, 2010) 48x60 mm No. 1 cover slip, four square 125-μm thick mylar shims serving as spacers, and an untreated 22x22 mm No. 1 cover slip, secured together with UV-curable adhesive. First, 0.8% anti-tubulin antibody (BioRad YL1/2) in ice-cold BRB-80 buffer (80 mM PIPES pH = 6.9, 1 mM MgCl_2_, 1 mM EGTA) is incubated in the flow cell, followed by 0.2 mg/ml NEM myosin in ice-cold Myosin Buffer (25 mM Imidazole pH = 7.4, 300 mM KCl, 4 mM EGTA, 4 mM MgCl_2_) for an additional 5 minutes. Both flow channels are washed with Wash Buffer (Actin Buffer plus 20 μM Taxol and 10 mM DTT) and then blocked with Block Buffer (BRB-80 plus 5% w/v Pluronic F-127) for 2 minutes. After washing with Wash Buffer, fluorescent MTs and actin are added sequentially with their order of addition determining which filament is on top in the intersection (Ross *et al*., 2008b; Schroeder *et al*., 2012; Lombardo *et al*., 2017; Bensel *et al*., 2024). MTs used are a 1:300 dilution of rhodamine-labelled MTs prepared as described previously (Bensel *et al*., 2024, 2025) into BRB-80 with 20 μM Taxol, while actin is a 1:30 dilution into Actin Buffer of a 1 μM stock of actin labelled with Alexa-fluor 488 phalloidin. Either MTs or actin were perfused into one channel, incubated for 5 minutes, and washed with Wash Buffer. Then the other filament type was perfused into the perpendicular channel, incubated for 5 minutes, and both channels were washed with Wash Buffer. Finally, liposomes with bound MyoVa, K543, and melanophilin (except for no melanophilin control) were diluted to a final concentration of ∼160 pM in Motility Buffer (Actin Buffer with with 2 mM MgATP, 20 μM paclitaxel, 0.5 mg/mL Casein, 0.5% w/v Pluronic F-127, 5 mM creatine phosphate, 0.4 mg/mL creatine phosphokinase, 10 mM DTT, 3.5 mg/mL glucose, 40 μg/mL glucose oxidase, and 27 μg/mL catalase) and perfused into the flow cell. Liposome motility was imaged on a custom-built TIRF microscope (described previously, (Lombardo *et al*., 2017; Bensel *et al*., 2024)) taking a 5-minute video at 10 frames per second in each ROI.

#### Data analysis

Data were analyzed both by visually assessing videos and by kymography (Schindelin *et al*., 2012), as described previously (Bensel *et al*., 2024, 2025). Analysis was restricted to liposomes which entered the intersection on the bottom track, where liposome track choice could be determined directly from the video. Pause lifetimes in the intersection were measured by generating a kymograph for the actin track and the MT track, pseudo-coloring the kymographs cyan for actin and yellow for MT, and overlaying the kymographs aligned to the intersection point. In this case, a vertical line present in both channels at the intersection point indicates a liposome that paused in the intersection. The shortest pause that could be identified confidently was 3 frames (0.3 s), so shorter pauses are excluded from analysis. Error bars for intersection outcomes were determined by bootstrapping analysis in R (R Core Team, 2025), and pause lifetime statistical analysis was performed in R.

### Melanophilin binding assay

Melanophilin binding assays were performed by first labeling melanophilin with quantum dots (Qdot 655 Streptavidin Conjugate, ThermoFisher) at a 4-fold molar excess to melanophilin. To ensure complete labeling, the quantum dot – melanophilin mixture was incubated for 1 hour on ice. During this time, flow cells were prepared using a 24x60 mm silanized (Dixit and Ross, 2010) No. 1 cover slip, mylar shims, and a 22x22 mm No. 1 cover slip, secured together with UV-curable adhesive. The cover slips were coated first with 0.8% w/v anti-tubulin antibody in BRB-80 for 5 minutes, washed with Wash Buffer, coated with 0.2 mg/ml NEM myosin for 5 minutes and washed again. Flow cells are blocked with Block Buffer for 2 minutes and washed prior to sequential addition of 1:300 rhodamine MTs in BRB-80 plus 20 μM Taxol for 5 minutes and 1:30 Alexa-fluor 488 labeled actin in Actin Buffer for 5 minutes followed by a wash with Wash Buffer. Quantum dot labelled melanophilin was diluted into Motility Buffer at a final concentration of 1 nm melanophilin/4 nm quantum dots and introduced into the flow cell. Melanophilin binding was imaged by TIRF microscopy on a custom-built microscope, taking a 1-minute video at 10 frames per second at each ROI.

#### Data analysis

Melanophilin binding density was quantified by kymography. Kymographs were drawn on actin and MT tracks and reproduced in the Qdot channel (Fig. 3E). For each kymograph, the length of the track, number of statically bound particles, and number of diffusing particles were recorded. Therefore for each analyzed track, a melanophilin binding density (bound particles per μm track length) and diffusive fraction were determined and plotted as dot plots per condition (Fig. 3F, G). Statistical comparisons of binding density and diffusive fraction were performed in R. Melanophilin diffusion constants on actin and MTs were measured using a custom pipeline in Fiji and R. For each binding assay video, the images of the actin and MTs were separately thresholded and converted to skeletons using the ‘Skeletonize 2D/3D’ plugin, prior to exporting these binary skeletons as .csv files. Visible particles in the Qdot channel were localized and linked into tracks using the DoM plugin (Katrukha *et al*., 2022) and exported as .csv files. The skeleton and Qdot tracking files were imported into R, where Qdot traces were first filtered such that only trajectories with at least 20 tracked points were considered, to eliminate brief, nonspecific interactions. Each remaining trace was referenced to the XY-coordinates of the actin and MT skeletons to determine, using a KD-tree approach (Bentley, 1975; Arya *et al*., 1998; Jefferis *et al*., 2013), if the trace was within a user-defined distance of an actin or MT. Traces that were within this distance threshold of either no track or both an actin and MT were discarded. Then the (X,Y,t) coordinates of the remaining traces were converted to a tangential distance along the assigned track, and this tangential distance over time was used to compute an independent diffusion coefficient (*D*) for each individual Qdot trace based on the Mean Squared Displacement (*MSD*) of the trace with a maximum lag of 10 frames. Individual diffusion constants for each particle were partitioned by melanophilin state and the assigned track, then plotted as Cumulative Distribution plots in Fig. 3H, with median *D* values reported, and statistical analysis performed in R. The equation used to compute *D* was:

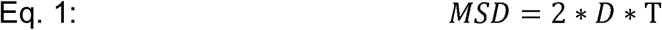

#### Liposome motility assay

Liposomes were prepared by incubating either K543 or MyoVa HMM at a 5:1 molar excess to liposomes as a control, with the addition of untreated, phosphorylated, or dephosphorylated melanophilin at a 5:1 molar excess to the liposomes. The mixture of liposomes, motor, and melanophilin was incubated on ice for 1 hour to allow for the binding reaction to complete. Flow cells were prepared by affixing a 22x22 mm No. 1 cover slip to a silanized 24x60 mm No. 1 cover slip with mylar shims and UV curable adhesive. The surface of the flow cell was then coated with either 0.2 mg/ml NEM myosin for actin-based motility or 0.8% w/v anti-tubulin antibody (BioRad YL1/2) for MT-based motility and incubated for 5 minutes prior to a wash with Wash Buffer. The flow cell was then blocked with 5% Pluronic F-127 in BRB-80 for 2 minutes and again washed. Either 1:30 AlexaFluor-488 actin or 1:300 rhodamine MTs were introduced to the flow cell and incubated for 5 minutes prior to a final wash. Liposomes, prepared as described above, were diluted to a final concentration of ∼160 pM in Motility Buffer and introduced into the flow cell. Motility was imaged by TIRF microscopy on a custom-built microscope, taking a 5-minute video at 10 frames per second at each ROI.

#### Data analysis

Liposome motility was analyzed by kymography as described previously (Bensel *et al*., 2024). Briefly, a kymograph was drawn along the track of interest and reproduced in the liposome channel. In the kymograph, time is shown on the vertical axis while distance is shown on the horizontal axis. A moving liposome will be observed as a slanted line on the kymograph. Each moving liposome is analyzed by drawing a bounding box around liposome trajectory, where the width of the box gives the run length and the velocity is defined as the width of the box divided by the height of the box. Whether or not the liposome reached the end of the track without detaching is recorded, and liposome runs that reach the end of the track are considered ‘end events.’ Furthermore, the length of each analyzed filament, i.e., the width of the full kymograph, is recorded, and used to derive a filament length distribution for each experimental condition (Fig. 4D and H, insets). Run length and velocity data are displayed as dot plots with an overlaid box plot, and end event frequencies are plotted separately as bar graphs (see Fig. 4). Statistical analysis to compare distributions was performed in R.

## Author contributions

B.M.B, K.M.T., and D.M.W designed research; B.M.B., P.M.F., J.E.M., O.G., and M.J.P. performed research, B.M.B., M.J.P., K.M.T., and D.M.W. analyzed data, and B.M.B., M.J.P., K.M.T., and D.M.W. wrote the paper.

## Competing interests

The authors declare no competing interests.

## Acknowledgements

We would like to acknowledge Shane R. Nelson for supporting in developing data analysis scripts, Guy Kennedy for TIRF microscopy support and training. We would also like to thank current and former members of the Warshaw and Trybus labs for their invaluable input, discussions, and support. The work published here would not have been possible without the contributions of those who have played a role in the creation, distribution, and maintenance of the open-source software packages used in this study, particularly r-project.org and ImageJ.nih.gov. Research reported in this publication was supported by the National Institutes of Health under Award Numbers T32HL076122 (to B.M.B.), F32GM140618 (to B.M.B.), R35GM141743 (to D.M.W.), R35GM136288 (to K.M.T.), and R01HL157487 (to M.J.P.). The content is solely the responsibility of the authors and does not necessarily represent the official views of the National Institutes of Health. We also thank Arnold and Mariel Goran for their generous gift to D.M.W.

## Abbreviations

(MT): microtubule
(HMM): heavy meromyosin
(PKA): protein kinase A

## Notes

### Competing Interest Statement

The authors have declared no competing interest.

### Summary of Updates

Editorial changes to better contextualize the study with existing literature.

## References

Arya, S, Mount, DM, Netanyahu, NS, Silverman, R, and Wu, AY (1998). An optimal algorithm for approximate nearest neighbor searching fixed dimensions. J ACM 45, 891–923.

Bao, M, Gempeler, M, and Campiche, R (2025). Melanosome Transport and Processing in Skin Pigmentation: Mechanisms and Targets for Pigmentation Modulation. IJMS 26, 8630.

Bensel, BM, Previs, SB, Bookwalter, C, Trybus, KM, Walcott, S, and Warshaw, DM (2024). Kinesin-1-transported liposomes prefer to go straight in 3D microtubule intersections by a mechanism shared by other molecular motors. Proc Natl Acad Sci USA 121, e2407330121.

Bensel, BM, Previs, SB, Fagnant, PM, Trybus, KM, Walcott, S, and Warshaw, DM (2025). Kinesin-1 autoinhibition tunes cargo transport by motor ensembles. Biophysical Journal 124, 3278–3290.

Bentley, JL (1975). Multidimensional binary search trees used for associative searching. Commun ACM 18, 509–517.

Bentley, M, and Banker, G (2016). The cellular mechanisms that maintain neuronal polarity. Nat Rev Neurosci 17, 611–622.

Bodakuntla, S, Jijumon, AS, Villablanca, C, Gonzalez-Billault, C, and Janke, C (2019). Microtubule-Associated Proteins: Structuring the Cytoskeleton. Trends in Cell Biology 29, 804–819.

Brozzi, F, Diraison, F, Lajus, S, Rajatileka, S, Philips, T, Regazzi, R, Fukuda, M, Verkade, P, Molnár, E, and Váradi, A (2012). Molecular Mechanism of Myosin Va Recruitment to Dense Core Secretory Granules. Traffic 13, 54–69.

Cai, D, Hoppe, AD, Swanson, JA, and Verhey, KJ (2007). Kinesin-1 structural organization and conformational changes revealed by FRET stoichiometry in live cells. The Journal of Cell Biology 176, 51–63.

Carrier, M, Joint, M, Lutzing, R, Page, A, and Rochette-Egly, C (2016). Phosphoproteome and Transcriptome of RA-Responsive and RA-Resistant Breast Cancer Cell Lines. PLoS ONE 11, e0157290.

Carrington, G, Fatima, U, Caramujo, I, Lewis, T, Casas-Mao, D, and Peckham, M (2024). A multiscale approach reveals the molecular architecture of the autoinhibited kinesin KIF5A. J Biol Chem 300, 105713.

Chaudhary, AR, Berger, F, Berger, CL, and Hendricks, AG (2018). Tau directs intracellular trafficking by regulating the forces exerted by kinesin and dynein teams. Traffic 19, 111–121.

Coppola, T, Frantz, C, Perret-Menoud, V, Gattesco, S, Hirling, H, and Regazzi, R (2002). Pancreatic β-Cell Protein Granuphilin Binds Rab3 and Munc-18 and Controls Exocytosis. MBoC 13, 1906–1915.

Deeb, SK, Guzik-Lendrum, S, Jeffrey, JD, and Gilbert, SP (2019). The ability of the kinesin-2 heterodimer KIF3AC to navigate microtubule networks is provided by the KIF3A motor domain. Journal of Biological Chemistry 294, 20070–20083.

Derr, ND, Goodman, BS, Jungmann, R, Leschziner, AE, Shih, WM, and Reck-Peterson, SL (2012). Tug-of-War in Motor Protein Ensembles Revealed with a Programmable DNA Origami Scaffold. Science 338, 662–665.

Desnos, C, Huet, S, and Darchen, F (2007a). ‘Should I stay or should I go?’: myosin V function in organelle trafficking. Biology of the Cell 99, 411–423.

Desnos, C, Huet, S, Fanget, I, Chapuis, C, Böttiger, C, Racine, V, Sibarita, J-B, Henry, J-P, and Darchen, F (2007b). Myosin Va Mediates Docking of Secretory Granules at the Plasma Membrane. J Neurosci 27, 10636–10645.

Desnos, C, Schonn, J-S, Huet, S, Tran, VS, El-Amraoui, A, Raposo, G, Fanget, I, Chapuis, C, Ménasché, G, De Saint Basile, G, et al. (2003). Rab27A and its effector MyRIP link secretory granules to F-actin and control their motion towards release sites. The Journal of Cell Biology 163, 559–570.

Dixit, R, and Ross, JL (2010). Studying Plus-End Tracking at Single Molecule Resolution Using TIRF Microscopy. In: Methods in Cell Biology, Elsevier, 543–554.

Dogterom, M, and Koenderink, GH (2019). Actin–microtubule crosstalk in cell biology. Nat Rev Mol Cell Biol 20, 38–54.

Fan, R, and Lai, K (2022). Understanding how kinesin motor proteins regulate postsynaptic function in neuron. The FEBS Journal 289, 2128–2144.

Feng, Q, Mickolajczyk, KJ, Chen, G-Y, and Hancock, WO (2018). Motor Reattachment Kinetics Play a Dominant Role in Multimotor-Driven Cargo Transport. Biophysical Journal 114, 400–409.

Fourriere, L, Jimenez, AJ, Perez, F, and Boncompain, G (2020). The role of microtubules in secretory protein transport. Journal of Cell Science 133, jcs237016.

Fukuda, M, and Kuroda, TS (2002). Slac2-c (Synaptotagmin-like Protein HomologueLacking C2 Domains-c), a Novel Linker Protein that Interacts with Rab27, Myosin Va/VIIa, and Actin. Journal of Biological Chemistry 277, 43096–43103.

Furuta, K, Furuta, A, Toyoshima, YY, Amino, M, Oiwa, K, and Kojima, H (2013). Measuring collective transport by defined numbers of processive and nonprocessive kinesin motors. Proc Natl Acad Sci USA 110, 501–506.

Grover, R, Fischer, J, Schwarz, FW, Walter, WJ, Schwille, P, and Diez, S (2016). Transport efficiency of membrane-anchored kinesin-1 motors depends on motor density and diffusivity. Proc Natl Acad Sci USA 113.

Gunning, P, O’neill, G, and Hardeman, E (2008). Tropomyosin-Based Regulation of the Actin Cytoskeleton in Time and Space. Physiological Reviews 88, 1–35.

Hammer, JA, and Sellers, JR (2012). Walking to work: roles for class V myosins as cargo transporters. Nat Rev Mol Cell Biol 13, 13–26.

Hammer, JA, and Wu, XS (2002). Rabs grab motors: defining the connections between Rab GTPases and motor proteins. Current Opinion in Cell Biology 14, 69–75.

Heaslip, AT, Nelson, SR, Lombardo, AT, Beck Previs, S, Armstrong, J, and Warshaw, DM (2014). Cytoskeletal Dependence of Insulin Granule Movement Dynamics in INS-1 Beta-Cells in Response to Glucose. PLoS ONE 9, e109082.

Hodges, AR, Krementsova, EB, and Trybus, KM (2007). Engineering the Processive Run Length of Myosin V. Journal of Biological Chemistry 282, 27192–27197.

Jefferis, G, Kemp, SE, Arya, S, and Mount, D (2013). RANN: Fast Nearest Neighbour Search (Wraps ANN Library) Using L2 Metric.

Jenkins, B, Decker, H, Bentley, M, Luisi, J, and Banker, G (2012). A novel split kinesin assay identifies motor proteins that interact with distinct vesicle populations. Journal of Cell Biology 198, 749–761.

Katrukha, E, Jalmar Teeuw, Bmccloin, and Braber, JD (2022). ekatrukha/DoM_Utrecht: Detection of Molecules 1.2.5, Zenodo.

Kunwar, A, and Mogilner, A (2010). Robust transport by multiple motors with nonlinear force–velocity relations and stochastic load sharing. Phys Biol 7, 016012.

Lipka, J, Kapitein, LC, Jaworski, J, and Hoogenraad, CC (2016). Microtubule-binding protein doublecortin-like kinase 1 (DCLK1) guides kinesin-3-mediated cargo transport to dendrites. The EMBO Journal 35, 302–318.

Lombardo, AT, Nelson, SR, Ali, MY, Kennedy, GG, Trybus, KM, Walcott, S, and Warshaw, DM (2017). Myosin Va molecular motors manoeuvre liposome cargo through suspended actin filament intersections in vitro. Nat Commun 8, 15692.

Lundby, A, Secher, A, Lage, K, Nordsborg, NB, Dmytriyev, A, Lundby, C, and Olsen, JV (2012). Quantitative maps of protein phosphorylation sites across 14 different rat organs and tissues. Nat Commun 3, 876.

Mayr, MI, Storch, M, Howard, J, and Mayer, TU (2011). A Non-Motor Microtubule Binding Site Is Essential for the High Processivity and Mitotic Function of Kinesin-8 Kif18A. PLoS ONE 6, e27471.

McKenney, RJ, Huynh, W, Tanenbaum, ME, Bhabha, G, and Vale, RD (2014). Activation of cytoplasmic dynein motility by dynactin-cargo adapter complexes. Science 345, 337– 341.

Michalek, AJ, Kennedy, GG, Warshaw, DM, and Ali, MY (2015). Flexural Stiffness of Myosin Va Subdomains as Measured from Tethered Particle Motion. Journal of Biophysics 2015, 1–9.

Minoura, I, Katayama, E, Sekimoto, K, and Muto, E (2010). One-Dimensional Brownian Motion of Charged Nanoparticles along Microtubules: A Model System for Weak Binding Interactions. Biophysical Journal 98, 1589–1597.

Mizoguchi, A, Yano, Y, Hamaguchi, H, Yanagida, H, Ide, C, Zahraoui, A, Shirataki, H, Sasaki, T, and Takai, Y (1994). Localization of Rabphilin-3A on the Synaptic Vesicle. Biochemical and Biophysical Research Communications 202, 1235–1243.

Monroy, BY, Tan, TC, Oclaman, JM, Han, JS, Simó, S, Niwa, S, Nowakowski, DW, McKenney, RJ, and Ori-McKenney, KM (2020). A Combinatorial MAP Code Dictates Polarized Microtubule Transport. Developmental Cell 53, 60–72.e4.

Nascimento, AA, Roland, JT, and Gelfand, VI (2003). Pigment Cells: A Model for the Study of Organelle Transport. Annu Rev Cell Dev Biol 19, 469–491.

Nayak, CR, and Rutenberg, AD (2011). Quantification of Fluorophore Copy Number from Intrinsic Fluctuations during Fluorescence Photobleaching. Biophysical Journal 101, 2284–2293.

Nelson, PC, Zurla, C, Brogioli, D, Beausang, JF, Finzi, L, and Dunlap, D (2006). Tethered Particle Motion as a Diagnostic of DNA Tether Length. J Phys Chem B 110, 17260–17267.

Nelson, SR, Trybus, KM, and Warshaw, DM (2014). Motor coupling through lipid membranes enhances transport velocities for ensembles of myosin Va. Proc Natl Acad Sci USA 111.

Oberhofer, A, Reithmann, E, Spieler, P, Stepp, WL, Zimmermann, D, Schmid, B, Frey, E, and Ökten, Z (2020). Molecular underpinnings of cytoskeletal cross-talk. Proc Natl Acad Sci U S A 117, 3944–3952.

Oberhofer, A, Spieler, P, Rosenfeld, Y, Stepp, WL, Cleetus, A, Hume, AN, Mueller-Planitz, F, and Ökten, Z (2017). Myosin Va’s adaptor protein melanophilin enforces track selection on the microtubule and actin networks in vitro. Proc Natl Acad Sci U S A 114, E4714–E4723.

O’Leary, TS, Snyder, J, Sadayappan, S, Day, SM, and Previs, MJ (2019). MYBPC3 truncation mutations enhance actomyosin contractile mechanics in human hypertrophic cardiomyopathy. Journal of Molecular and Cellular Cardiology 127, 165–173.

Osunbayo, O, Butterfield, J, Bergman, J, Mershon, L, Rodionov, V, and Vershinin, M (2015). Cargo Transport at Microtubule Crossings: Evidence for Prolonged Tug-of-War between Kinesin Motors. Biophysical Journal 108, 1480–1483.

Pan, X, Cao, Y, Stucchi, R, Hooikaas, PJ, Portegies, S, Will, L, Martin, M, Akhmanova, A, Harterink, M, and Hoogenraad, CC (2019). MAP7D2 Localizes to the Proximal Axon and Locally Promotes Kinesin-1-Mediated Cargo Transport into the Axon. Cell Reports 26, 1988–1999.e6.

Pimm, ML, and Henty-Ridilla, JL (2021). New twists in actin–microtubule interactions. MBoC 32, 211–217.

Pouget, N, Dennis, C, Turlan, C, Grigoriev, M, Chandler, M, and Salomé, L (2004). Single-particle tracking for DNA tether length monitoring. Nucleic Acids Res 32, e73.

Previs, MJ, VanBuren, P, Begin, KJ, Vigoreaux, JO, LeWinter, MM, and Matthews, DE (2008). Quantification of Protein Phosphorylation by Liquid Chromatography−Mass Spectrometry. Anal Chem 80, 5864–5872.

R Core Team (2025). R: A language and environment for statistical computing., Vienna, Austria: R Foundation for Statistical Computing.

Reck-Peterson, SL, Redwine, WB, Vale, RD, and Carter, AP (2018). The cytoplasmic dynein transport machinery and its many cargoes. Nat Rev Mol Cell Biol 19, 382–398.

Rogers, SL, Tint, IS, Fanapour, PC, and Gelfand, VI (1997). Regulated bidirectional motility of melanophore pigment granules along microtubules *in vitro*. Proc Natl Acad Sci USA 94, 3720–3725.

Ross, JL, Ali, MY, and Warshaw, DM (2008a). Cargo transport: molecular motors navigate a complex cytoskeleton. Current Opinion in Cell Biology 20, 41–47.

Ross, JL, Shuman, H, Holzbaur, ELF, and Goldman, YE (2008b). Kinesin and Dynein-Dynactin at Intersecting Microtubules: Motor Density Affects Dynein Function. Biophysical Journal 94, 3115–3125.

Sarpangala, N, and Gopinathan, A (2022). Cargo surface fluidity can reduce inter-motor mechanical interference, promote load-sharing and enhance processivity in teams of molecular motors. PLoS Comput Biol 18, e1010217.

Schindelin, J, Arganda-Carreras, I, Frise, E, Kaynig, V, Longair, M, Pietzsch, T, Preibisch, S, Rueden, C, Saalfeld, S, Schmid, B, et al. (2012). Fiji: an open-source platform for biological-image analysis. Nat Methods 9, 676–682.

Schroeder, HW, Hendricks, AG, Ikeda, K, Shuman, H, Rodionov, V, Ikebe, M, Goldman, YE, and Holzbaur, ELF (2012). Force-Dependent Detachment of Kinesin-2 Biases Track Switching at Cytoskeletal Filament Intersections. Biophysical Journal 103, 48–58.

Sckolnick, M, Krementsova, EB, Warshaw, DM, and Trybus, KM (2013). More than just a cargo adapter, melanophilin prolongs and slows processive runs of myosin Va. J Biol Chem 288, 29313–29322.

Sckolnick, M, Krementsova, EB, Warshaw, DM, and Trybus, KM (2016). Tropomyosin isoforms bias actin track selection by vertebrate myosin Va. MBoC 27, 2889–2897.

Seabra, MC, and Coudrier, E (2004). Rab GTPases and Myosin Motors in Organelle Motility. Traffic 5, 393–399.

Sheets, L, Ransom, DG, Mellgren, EM, Johnson, SL, and Schnapp, BJ (2007). Zebrafish Melanophilin Facilitates Melanosome Dispersion by Regulating Dynein. Current Biology 17, 1721–1734.

Sköld, HN, Norström, E, and Wallin, M (2002). Regulatory Control of Both Microtubule- and Actin-dependent Fish Melanosome Movement. Pigment Cell Research 15, 357– 366.

Soppina, V, Norris, SR, Dizaji, AS, Kortus, M, Veatch, S, Peckham, M, and Verhey, KJ (2014). Dimerization of mammalian kinesin-3 motors results in superprocessive motion. Proc Natl Acad Sci USA 111, 5562–5567.

Spiliotis, ET, and Nakos, K (2021). Cellular functions of actin- and microtubule-associated septins. Current Biology 31, R651–R666.

Stern, JL, Lessard, DV, Hoeprich, GJ, Morfini, GA, and Berger, CL (2017). Phosphoregulation of Tau modulates inhibition of kinesin-1 motility. MBoC 28, 1079– 1087.

Tan, Z, Yue, Y, Leprevost, F, Haynes, S, Basrur, V, Nesvizhskii, AI, Verhey, KJ, and Cianfrocco, MA (2023). Autoinhibited kinesin-1 adopts a hierarchical folding pattern. eLife 12, RP86776.

Tjioe, M, Shukla, S, Vaidya, R, Troitskaia, A, Bookwalter, CS, Trybus, KM, Chemla, YR, and Selvin, PR (2019). Multiple kinesins induce tension for smooth cargo transport. eLife 8, e50974.

Trybus, KM (2008). Myosin V from head to tail. Cell Mol Life Sci 65, 1378–1389.

Vale, RD (2003). The Molecular Motor Toolbox for Intracellular Transport. Cell 112, 467– 480.

Varadi, A, Ainscow, EK, Allan, VJ, and Rutter, GA (2002). Involvement of conventional kinesin in glucose-stimulated secretory granule movements and exocytosis in clonal pancreatic β-cells. Journal of Cell Science 115, 4177–4189.

Varadi, A, Tsuboi, T, and Rutter, GA (2005). Myosin Va Transports Dense Core Secretory Vesicles in Pancreatic MIN6 β-Cells. MBoC 16, 2670–2680.

Verdeny-Vilanova, I, Wehnekamp, F, Mohan, N, Álvarez, ÁS, Borbely, JS, Otterstrom, JJ, Lamb, DC, and Lakadamyali, M (2017). 3D motion of vesicles along microtubules helps them to circumvent obstacles in cells. Journal of Cell Science, jcs.201178.

Verhey, KJ, and Hammond, JW (2009). Traffic control: regulation of kinesin motors. Nat Rev Mol Cell Biol 10, 765–777.

Vershinin, M, Carter, BC, Razafsky, DS, King, SJ, and Gross, SP (2007). Multiple-motor based transport and its regulation by Tau. Proc Natl Acad Sci USA 104, 87–92.

Wong, S, and Weisman, LS (2021). Roles and regulation of myosin V interaction with cargo. Advances in Biological Regulation 79, 100787.

Wood, NB, Kelly, CM, O’Leary, TS, Martin, JL, and Previs, MJ (2022). Cardiac Myosin Filaments are Maintained by Stochastic Protein Replacement. Molecular & Cellular Proteomics 21, 100274.

Wu, X, Bowers, B, Rao, K, Wei, Q, and Hammer, JA (1998). Visualization of Melanosome Dynamics within Wild-Type and Dilute Melanocytes Suggests a Paradigm for Myosin V Function In Vivo. The Journal of Cell Biology 143, 1899–1918.

Wu, XS, Rao, K, Zhang, H, Wang, F, Sellers, JR, Matesic, LE, Copeland, NG, Jenkins, NA, and Hammer, JA (2002). Identification of an organelle receptor for myosin-Va. Nat Cell Biol 4, 271–278.

Wu, XS, Tsan, GL, and Hammer, JA (2005). Melanophilin and myosin Va track the microtubule plus end on EB1. J Cell Biol 171, 201–207.

Zanivan, S, Gnad, F, Wickström, SA, Geiger, T, Macek, B, Cox, J, Fässler, R, and Mann, M (2008). Solid Tumor Proteome and Phosphoproteome Analysis by High Resolution Mass Spectrometry. J Proteome Res 7, 5314–5326.

