## Supplementary Materials for "The Phosphorylation State of Melanophilin, a Myosin Va Adapter Protein, Biases the Tug-of-War Between Myosin Va and Kinesin-1 on Liposomes at Actin-Microtubule Intersections *In Vitro*"

Supplementary Materials for “Melanophilin, a Myosin Va Adapter Protein, Biases Track Selection of Myosin Va-and Kinesin-1-Transported Liposomes at Actin-Microtubule Intersections *In Vitro*”

Supplementary Movies

Movie S1: Liposomes starting on actin can exit on actin in actin-MT intersections. A liposome (magenta) traverses an actin-MT intersection starting on actin (cyan), with the intersection geometry illustrated in Fig. 2A. After reaching the intersection, the liposome pauses at the MT (yellow) and then exits on the actin filament it entered on. The liposomes is decorated with 5 K543, 5 MyoVa HMM, and 5 PKA-treated melanophilin. Scale bar = 2 μm, video collected at 10 frames per second, video playback at 2X real time.

Movie S2: Liposomes starting on actin can exit on the MT in actin-MT intersections. A liposome (magenta) traverses an actin-MT intersection starting on actin (cyan), with the intersection geometry illustrated in Fig. 2A. After reaching the intersection, the liposome pauses and then exits on the MT (yellow). The liposome shown here is a control liposomes without melanophilin, decorated with 5 K543 and 5 MyoVa HMM. Scale bar = 2 μm, video collected at 10 frames per second, video playback at 2X real time.

Movie S3: Liposomes starting on a MT can exit on the MT in an actin-MT intersection. A liposome (magenta) traverses an actin (cyan)-MT (yellow) intersection starting on the MT, with the intersection geometry illustrated in Fig. 2E. The liposome reaches the intersection and exits on the MT without a noticeable pause. The liposome shown here is a control liposome without melanophilin and decorated with 5 K543 and 5 MyoVa HMM. Scale bar = 2 μm, video collected at 10 frames per second, video playback at 2X real time.

Movie S4: Liposomes starting on a MT can exit on the actin in an actin-MT intersection. A liposome (magenta) traverses an actin-MT intersection starting on the MT (yellow), with the intersection geometry illustrated in Fig. 2E. The liposome reaches the intersection, pauses, and then exits on the actin (cyan). The liposome shown here is a control without melanophilin and decorated with 5 K543 and 5 MyoVa HMM. Scale bar = 2 μm, video collected at 10 frames per second, video playback at 2X real time.

Movie S5: Melanophilin binding assay with PKA-treated melanophilin. Qdot-labeled, PKA-treated melanophilin (magenta) associates with actin (cyan) and MTs (yellow) *in vitro.* Melanophilin displays diffusive motion on MTs, but appears stationary on actin. Scale bar = 5 μm, video collected at 10 frames per second, video playback at 1.5X real time.

Movie S6: Melanophilin binding assay with untreated melanophilin. Qdot-labeled, untreated melanophilin (magenta) associates with actin (cyan) and MTs (yellow) *in vitro.* Melanophilin displays diffusive motion on MTs, but appears stationary on actin. Scale bar = 5 μm, video collected at 10 frames per second, video playback at 1.5X real time.

Movie S7: Melanophilin binding assay with dephosphorylated melanophilin. Qdot-labeled, dephosphorylated melanophilin (magenta) associates with actin (cyan) and MTs (yellow) *in vitro.* Melanophilin displays diffusive motion on MTs, but appears stationary on actin. Scale bar = 5 μm, video collected at 10 frames per second, video playback at 1.5X real time.

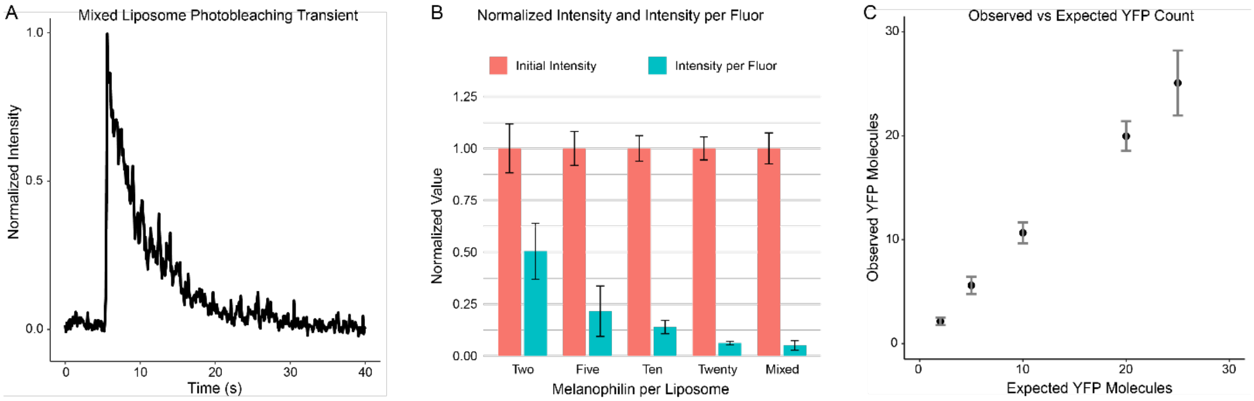


Figure S1. Photobleaching estimation of liposome-bound YFP-melanophilin molecules. A) Example photobleaching transient recorded for a liposome that was incubated with a 5-fold excess each of YFP-K543, YFP-MyoVa HMM, and YFP-melanophilin. This is expected to yield 25 liposome-bound YFP molecules – 2 each per K543 and MyoVa HMM dimers, and 1 per melanophilin. B) Bar graph of median ± SEM normalized initial intensity and normalized intensity per fluor for each incubation ratio used to derive a melanophilin standard binding curve, as well as mixed liposomes prepared as in A. C) Plot of observed YFP molecules versus expected YFP molecules corresponding to the incubation conditions shown in B, i.e., a melanophilin standard binding curve. The rightmost point, with an X-value of 25, corresponds to the mixed liposomes described in A.
